# *Draft genome sequence of Microbacterium fakhimi* sp. nov., a novel bacterium associated with the alga *Chlamydomonas reinhardtii*

**DOI:** 10.1101/2023.05.04.539371

**Authors:** Neda Fakhimi, María Jesus Torres, Jesús Delgado-Luque, Emilio Fernandez, Aurora Galván, Alexandra Dubini, David González-Ballester

**Author notes:** Corresponding author: Alexandra Dubini. Emails: Neda Fakhimi; M. J. Torres; J. Delgado; E. Fernández; A. Galván; A. Dubini; D. González-Ballester (.

## Abstract

*Microbacterium fakhimi* sp. nov. has been isolated from a contaminated algal culture (*Chlamydomonas reinhardtii*). Its genome has been fully sequenced (3,753,259 base pairs) and a tentative annotation is provided (3,704 genes). Both, genome information and growth tests suggest that *M. fakhimi* sp. nov. is auxotroph for biotin and thiamine and unable to use sulfate as sulfur (S) source. S-reduced forms, such as methionine and cysteine can support *M. fakhimi* sp. nov. growth. The potential biotechnological interest of this bacteria is discussed here and in a related research paper (Fakhimi et al., 2023b).

## INTRODUCTION

Microbacterium genus belongs to the Microbacteriaceae family (Order: Micrococcales; Class: Actinomycetia; Phylum: Actinomycetota). Microbacterium sp. are described as gram-positive, aerobic, non-spore-forming, rod-shaped and high G+C content bacteria. First identified by Orla-Jensen in 1919, Microbacterium are now composed of 129 validated species (https://lpsn.dsmz.de/genus/microbacterium) (Parte et al., 2020). One of the Microbacteriaceae family specificities that differentiated them from other Actinobacteria is their atypical peptidoglycan cell wall and their unsaturated respiratory menaquinones (Evtushenko and Takeuchi, 2006).

Microbacterium spp. are widely distributed and has been isolated from a broad range of habitats such as soils (Takeuchi and Hatano, 1998; Lu et al., 2019), deserts (Mandakovic et al., 2015; Yang et al., 2018; Mandakovic et al., 2020; Zhu et al., 2021), arctic zones (Reis-Mansur et al., 2019), waters (Yi et al., 2021), heavy metal and other contaminated ecosystems (Qu et al., 2013; Learman et al., 2019; Lu et al., 2019; Corretto et al., 2020; Martínez-Rodríguez et al., 2020; Mishra et al., 2021), dairy products (Vithanage et al., 2016; Bellassi et al., 2020; Bellassi et al., 2021), phyllosphere (Behrendt et al., 2001; Marquez-Santacruz et al., 2010a; Bano et al., 2022), algae and cyanobacteria cultures (Jones et al., 2016; Mitra et al., 2021; Zhou et al., 2021), and animal host (Gneiding et al., 2008; Amano et al., 2019; Heo et al., 2020). Interestingly, Microbacterium also appears to be a common laboratory contaminant (Salter et al., 2014). Thereby they are easy to isolate from many environments and may explain the in-depth study of this family. In fact, 30 new species have been discovered in the last 5 years.

Many Microbacterium spp. have been study due to their biotechnological interest in facilitating bioremediation (Qu et al., 2013; Corretto et al., 2015; Avramov et al., 2016; Learman et al., 2019; Lu et al., 2019; Ortet et al., 2019; Corretto et al., 2020; Heidari et al., 2020; Martínez-Rodríguez et al., 2020; Mishra et al., 2021; Mitra et al., 2021) or promoting plant growth (Vílchez et al., 2018; Singh and Singh, 2019; García-Fontana et al., 2020; Yang et al., 2021). Moreover, many *Microbacterium* spp. can produce several secondary metabolites with diverse biological activities. Those include terpenoids, siderophores-like compounds (Corretto et al., 2020), carotenoids like decaprenoxanthin (Xie et al., 2021), and isoflavones and diketopiperazines with antifungal activities (Savi et al., 2019).

Here we report the genome of a new Microbacterium specie isolated from a microalgae culture (*Chlamydomonas reinhardtii*). This new bacterium has been named as *Microbacterium fakhimi* sp. nov. Tentative gene annotation has established that *M. fakhimi sp*. nov. is lacking biotin and thiamine biosynthetic pathways as well as a functional sulfate assimilation pathway. Growth tests have confirmed the dependence for the sulfur(S)-containing metabolites biotin, thiamine, methionine, and cysteine.

The metabolic interactions established between *M. fakhimi* sp. nov. and *C. reinhardtii* are analyzed and discussed in a related publication (Fakhimi et al., 2023b) where we highlighted the ability of the M. *fakhimi-C. reinhardtii* consortium to mutualistically grow on media with mannitol, and to sustain H_2_ production.

## TECHNICAL PROCEDURES

### Isolation of *Microbacterium fakhimi* sp. nov

*M. fakhimi* sp. nov. was isolated from a fortuitously contaminated *Chlamydomonas reinhardtii* culture. Contamination occurred within the laboratory facilities (located at Campus Universitario de Rabanales, Cordoba, Spain). Initially, the algal *Chlamydomonas reinhardtii* culture was simultaneously contaminated with three different bacteria (Fakhimi et al., 2023b). Individual members of this bacterial community were isolated by sequential rounds of plate streaking in Yeast Extract Mannitol (YEM) medium, until 3 different types of bacterial colonies were visually identified. Colonies were grown separately, and the subsequent isolated DNA was used for PCR-amplification of their partial RNA 16S sequences. After sequencing, the three independently isolated bacteria were identified as members of the genus *Microbacterium, Stenotrophomonas, and Bacillus*. Their whole genomes sequences were obtained and further identified as *Microbacterium fakhimi* sp. nov., *Stenotrophomonas goyi* sp. nov. (Fakhimi et al., 2023b) and *Bacillus cereus*.

### Genome sequencing and assembling of *M. fakhimi* sp. nov

DNA isolation and whole genome sequencing using PacBio (Pacific Biosciences) were performed by SNPsaurus LLC (https://www.snpsaurus.com/). Whole genome sequencing generated 138,141 reads yielding 1,097,358,991 bases for 257 read depth over the genome (**Table 1**). Genome was assembled by The Bioknowledge Lab (BK-L) Ltd. with Flye 2.9.1 (Lin et al., 2016), Canu 2.2 (Koren et al., 2017) and Prokka 1.14.6 (Seemann, 2014), yielding a 3,753,259 pb circular genome. The genome completeness was checked by Busco (Manni et al., 2021) and was 93.9% complete, with 93.9% of the genome single copy and 0.0% duplicated. Any other prokaryotic contamination was discarded using ContEst16S (Lee et al., 2017). Search with PHASTER (Arndt et al., 2016) revealed no intact prophages.

**Table 1.** Main de novo sequencing and assembly statistics according to RAST and genome features of *Microbacterium fakhimi* sp. nov. according to RAST and NCBI-PGAP servers.

| Genome Size (pb) | Fold-coverage | GC content (%) | Nº of contigs | Type | Plasmids |
| --- | --- | --- | --- | --- | --- |
| 3,753,259 | 257 | 67.9 | 1 | circular | No |
|  | CDS | tRNAs | rRNAs | tmRNAs |  |
| RAST | 3,646 | 46 | 6 | - |  |
| NCBI-PGAP | 3,580 | 52 | 6 | 1 |  |

### Phylogenetic analysis

Phylogenetic analyses were performed using the TYGS (Type (Strain) Genome Server) (https://tygs.dsmz.de/) at Leibniz Institute DSMZ (German Collection of Microorganisms and Cell Cultures GmbH) (Meier-Kolthoff and Göker, 2019; Meier-Kolthoff et al., 2022). Information on nomenclature, synonymy and associated taxonomic literature was provided by TYGS’s sister database, the List of Prokaryotic names with Standing in Nomenclature (LPSN, available at https://lpsn.dsmz.de). The TYGS analysis was subdivided into 4 steps: 1) <u>Determination of closely related type strains</u>: Determination of closest type strain genomes was done in two complementary ways: First, genome was compared against all type strain genomes available in the TYGS database via the MASH algorithm (Ondov et al., 2016), and the ten type strains with the smallest MASH distances were chosen. Second, an additional set of ten closely related type strains was determined via the 16S rDNA gene sequences. These were extracted from the genome using RNAmmer (Lagesen et al., 2007) and each sequence was subsequently BLASTed (Camacho et al., 2009) against the 16S rDNA gene sequence of each of the currently 17208 type strains available in the TYGS database. This was used as a proxy to find the best 50 matching type strains (according to the bitscore) for the genome under study and to subsequently calculate precise distances using the Genome BLAST Distance Phylogeny approach (GBDP) under the algorithm ‘coverage’ and distance formula d5 (Meier-Kolthoff et al., 2013). These distances were finally used to determine the 10 closest type strain genomes for each of the user genomes. 2) <u>Pairwise comparison of genome sequences</u>: For the phylogenomic inference, all pairwise comparisons among the set of genomes were conducted using GBDP and accurate intergenomic distances inferred under the algorithm ‘trimming’ and distance formula d_5_ (Meier-Kolthoff et al., 2013). 100 distance replicates were calculated each. Digital DDH values and confidence intervals were calculated using the recommended settings of the GGDC 3.0 (Meier-Kolthoff et al., 2013; Meier-Kolthoff et al., 2022). 3) <u>Phylogenetic inference</u>: The resulting intergenomic distances were used to infer a balanced minimum evolution tree with branch support via FASTME 2.1.6.1 including SPR postprocessing (Lefort et al., 2015). Branch support was inferred from 100 pseudo-bootstrap replicates each. The trees were rooted at the midpoint (Farris, 1972) and visualized with PhyD3 (Kreft et al., 2017). 4) <u>Type-based species and subspecies clustering</u>: The type-based species clustering using a 70% dDDH radius around each of the 12 type strains was done as previously described (Meier-Kolthoff and Göker, 2019). Subspecies clustering was done using a 79% dDDH threshold as previously introduced (Meier-Kolthoff et al., 2014). dDDH values were provided according to the following GBDP formulas: formula d0 (a.k.a. GGDC formula 1): length of all HSPs divided by total genome length; formula d4 (a.k.a. GGDC formula 2): sum of all identities found in HSPs divided by overall HSP length; formula d6 (a.k.a. GGDC formula 3): sum of all identities found in HSPs divided by total genome length. Note: Formula d4 is independent of genome length and is thus robust against the use of incomplete draft genomes.

### Annotation

The NCBI’s Prokaryotic Genome Annotation Pipeline (PGAP) annotation service and the RAST tool kit, RASTtk at The Genome Annotation Service (Overbeek et al., 2014; Brettin et al., 2015) were used.

### Growth test and media

All the bacterial precultures were grown on Yeast Extract Mannitol (YEM) or Lysogeny broth (LB) media. Many growth experiment used Mineral Medium (MM) (Harris, 2008) supplemented with different nutrient sources. Tris-Acetate-Phosphate (TAP) (Harris, 2008) was also used occasionally. More specific details for each experiment can be found in in the corresponding figure and table captions. Bacterium cultures were incubated at 24-28°C and under continuous agitation (130 rpm).

### Pigments extraction and spectrophotometric evaluation

*M. fakhimi* sp. nov. was grown under continuous illumination (Photosynthetic Photon Flux Density (PPFD) of 60-90 μmol photon·m^-2^·s^-1^), 24ºC and 140 rpm. Pigments extraction was performed according to (Mitra et al., 2021). For this, 1 mL of cells were treated with 5mL of 100% methanol during 1 h in the dark at 20ºC for carotenoid extraction. After centrifugation for 1 minute at 13,000 rpm, the supernatant containing the extracted pigments were scanned for absorption and different wavelengths (from 400nm to 800nm) using a Beckman Coulter Spectrophotometer DU.

### Data availability

*Microbacterium fakhimi* sp. nov. has been deposited in the Spanish Type Culture Collection (CECT) with accession number CECT30765. Genome sequence for *Microbacterium fakhimi* sp. nov. has been deposited as GenBank CP116871 at NCBI. *M. fakhimi* sp. nov. has been submitted for patenting (OEPM Submission #P202330306).

## RESULTS

### Identification of *Microbacterium fakhimi* sp. nov

A fortuitous contaminated *Chlamydomonas reinhardtii* culture (strain 704) was studied due to its enhanced hydrogen production capability. This algal culture turned out to be contaminated with three different bacterial strain (Fakhimi et al., 2023b), one of them consisting in a yellow-pigmented bacterium (**Supplemental Fig. 1**). This bacterium was isolated after several rounds of plate streaking in the TYM medium. First, partial PCR amplification and sequencing of the ribosomal 16S gene allowed the identification of this bacteria as a member of the *Microbacterium* genus. Afterwards, the whole genome sequence was obtained. Genome assembling provided one single circular contig of 3,753,259 pb (**Table 1**). No plasmids or extrachromosomal elements were identified.

The NCBI-PGAP service identified 3,580 CDS plus some other genomic features (**Table 1**). Similarly, using the RAST server, 3,704 genes (3,646 CDS+ 52 rRNAs and tRNAs) were identified (**Table 1**). Out of these 3,646 CDS identified by RAST, 844 of them were in subsystems. Tentative genome annotation derived from the RAST server is available in **Supplemental Table 1**.

Phylogenetic analyses were performed with both, the whole genome (**Fig. 1a**) and the inferred 16S rDNAs (**Fig. 1b**). Pairwise comparisons with the closest type strains genomes are shown in **Supplemental Table 2**. Additional information of these closest type strains is in **Supplemental Table 3**. These phylogenetic analyses revealed that the sequenced genome belonged to a new *Microbacterium* specie; all dDDH values (d0, d4 and d6) were below 70% (Meier-Kolthoff et al., 2013) (**Supplemental Table 2**). This new bacterium was designated as *Microbacterium fakhimi* sp. nov. The closest related bacteria in terms of whole genome and 16S rDNA similarities were *Microbacterium phyllosphaerae* DSM 13468 and *Microbacterium foliorum* DSM 12966, respectively (**Fig. 1**). *M. fakhimi* sp. nov. genome was deposited in the NCBI (GenBank CP116871).

**Fig. 1.**
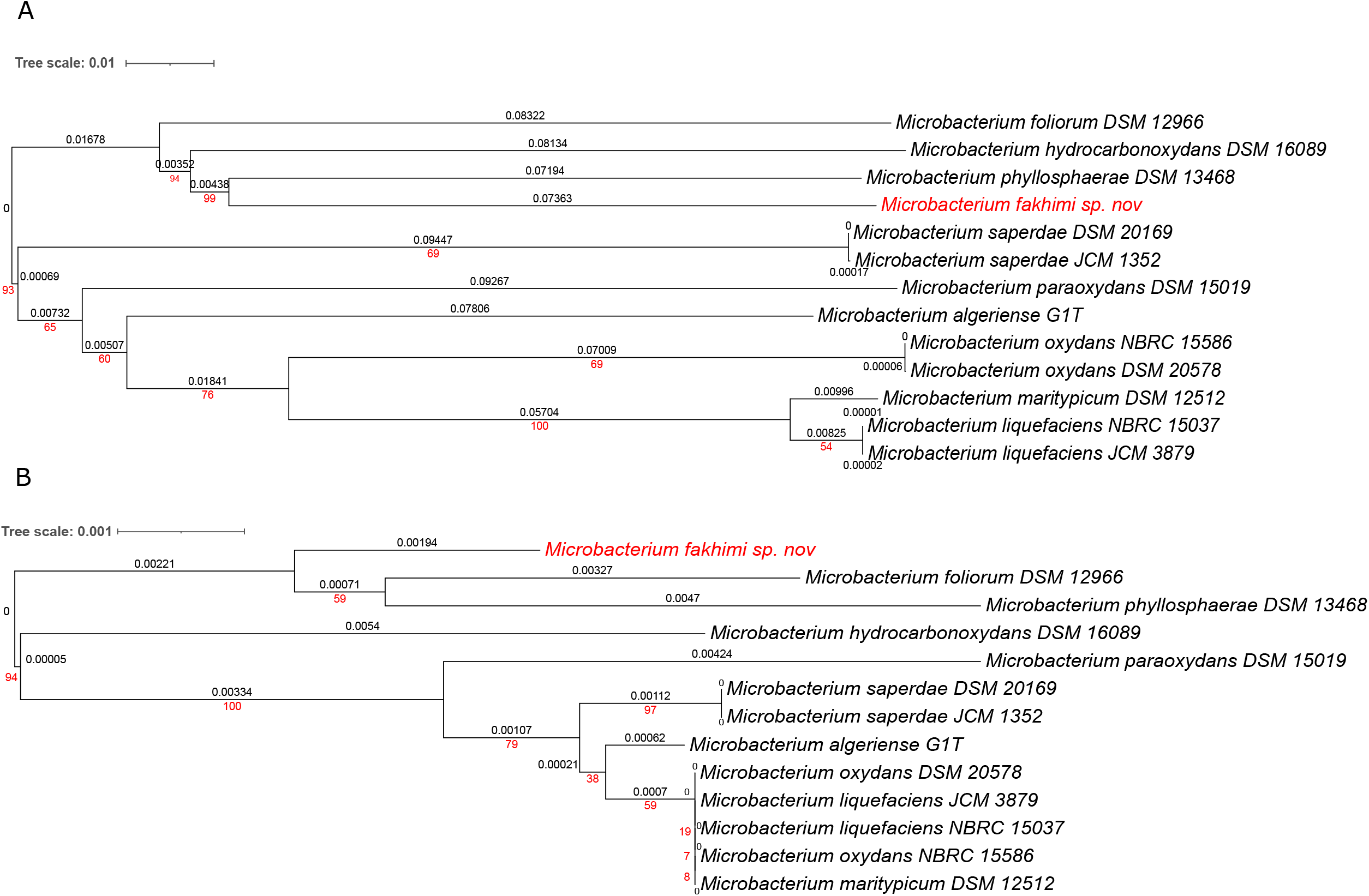
Phylogenetic trees for *Microbacterium fakhimi* and related closest bacteria based on genome (A) and 16S rDNA (B) Tree inferred with FastME 2.1.6.1 from GBDP distances calculated from genome and 16S rDNA gene sequences. The branch lengths are scaled in terms of GBDP distance formula d5. The numbers above branches are GBDP pseudo-bootstrap support values > 60 % from 100 replications, with an average branch support of 77.9 % (genomes) and 56.0 % (rDNA). The tree was rooted at the midpoint. Branch lengths (black) and bootstraps (red) values are indicated. Genome sizes: 3,314,049 - 4,323,590 pb; RNA16S lengths: 1,511 - 1,513 pb. Average δ statistics: 0.115 (genomes) and 0.092 (rDNAs) (Holland et al., 2002). Phylogenetic tree drawn with iTOL (Letunic and Bork, 2021).

BlastKOALA (Kanehisa et al., 2016) service allowed KEGG orthology assignments to characterize individual gene functions and reconstruct KEGG pathways of *M. fakhimi* genome (**Supplemental Table 4**). Some important pathways were either absent or incomplete in *M. fakhimi* sp. nov. including assimilation of sulfate (the whole assimilation pathway is missing), shikimate pathway (lack of shikimate dehydrogenase), and biosynthesis of biotin, thiamine, and cobalamin. On the other hand, putative complete pathways for the glyoxylate cycle and biosynthesis of riboflavin, pyridoxal-P, NAD, lipoic acid, coenzyme A and coenzyme F420, dTDP-L-rhamnose, UDP-N-acetyl-D-glucosamine, C5 and C10-C20 isoprenoids, among others, were present.

### Nutrient requirements of *M. fakhimi* sp. nov. to grow on sugars

In a recent publication, it was shown that *M. fakhimi* sp. nov. can grow on mannitol only if yeast extract was also supplemented (Fakhimi et al., 2023b). In the absent of yeast extract, cocultivation of *M. fakhimi* sp. nov. with the alga *Chlamydomonas reinhardtii* also allowed the bacterium growth.

To determine which components are present in the yeast extract, and potentially also in the algal broth medium, that allowed *M. fakhimi* sp. nov. to grow efficiently on mannitol, the nutrient requirements for this bacterium were studied.

First, it was observed that *M. fakhimi* sp. nov., did not grow in the MM medium supplemented with different carbon (C) sources (mannitol, glucose, or acetate), but required tryptone or yeast extract to grow. *M. fakhimi* sp. nov. even showed some growth when using albumin, indicating the capability of this bacterium to hydrolyze whole proteins (**Fig. 2**). Although tryptone, yeast extract or proteins could be used as nitrogen (N) and C sources by *M. fakhimi* sp. nov., they were likely mainly needed to complement other nutrient deficiencies. To investigate if these compounds were complementing any potential amino acid auxotrophy, the growth of the bacterium was tested with each individual amino acid supplementation (**Fig. 3a**). No single amino acid auxotrophy was observed. Still, when the 20 essential amino acids or pools of 19 different amino acids were used (each pool missing one single amino acid) the growth capability of *M. fakhimi* sp. nov. was restored (**Fig. 3b**). To solve if potential mixed amino acid auxotrophies were present in *M. fakhimi* sp. nov. different pools of amino acids were created (according to their biosynthetic similarities) (**Fig. 3c** and **5d**). As result, *M. fakhimi* sp. nov. partially restored the growth (relative to the media containing tryptone) when supplemented with cysteine or methionine, and especially when these two amino acids were added simultaneously **(Fig. 3d**). However, after several consecutive culturing rounds in cysteine- or methionine-containing media, no further growth was obtained (**Fig. 5**). Importantly, in the experiments depicted in Fig. 3, the precultures of *M. fakhimi* sp. nov. were done in the yeast-containing medium (LB medium). Hence, it was possible that, in addition to cysteine and methionine, other micronutrients originally present in the LB medium might be needed.

**Fig. 2.**
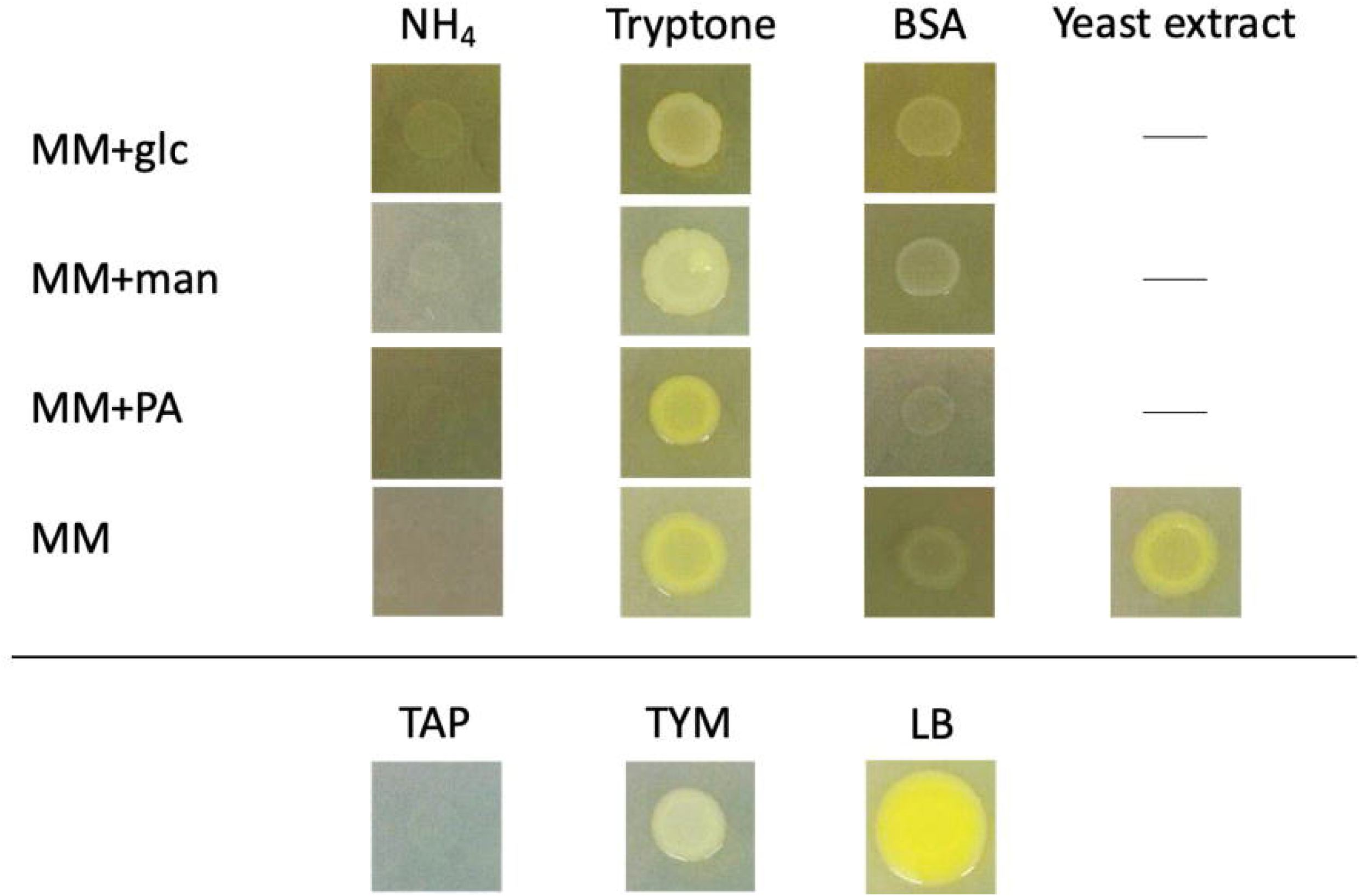
Growth (drop test) test for *M. fakhimi.* *M. fakhimi* was plated on MM supplemented with different C (5 g·L^-1^) and N (0.8 g·L^-1^) sources. C sources included glucose (glc), mannitol (man) and potassium acetate (PA), while N sources included tryptone (0.8 g·L^-1^), yeast extract (0.8 g·L^-1^) and non-hydrolyzed Bovine Serum Albumin (BSA)(0.5% w/v). Note that standard MM medium already contains 8 mM of ammonium. Bacterium precultures were harvested in logarithmic growth phase by centrifugation (8.000 g for 3 min), washed 3 times with MM and resuspended in the corresponding medium to a final OD of 0,02. Drops (6μL) of bacterium were added to the plates. Standard TAP, TYM and LB media were used as control media. Plates were incubated for 7 days at 25 ºC.

**Fig. 3.**
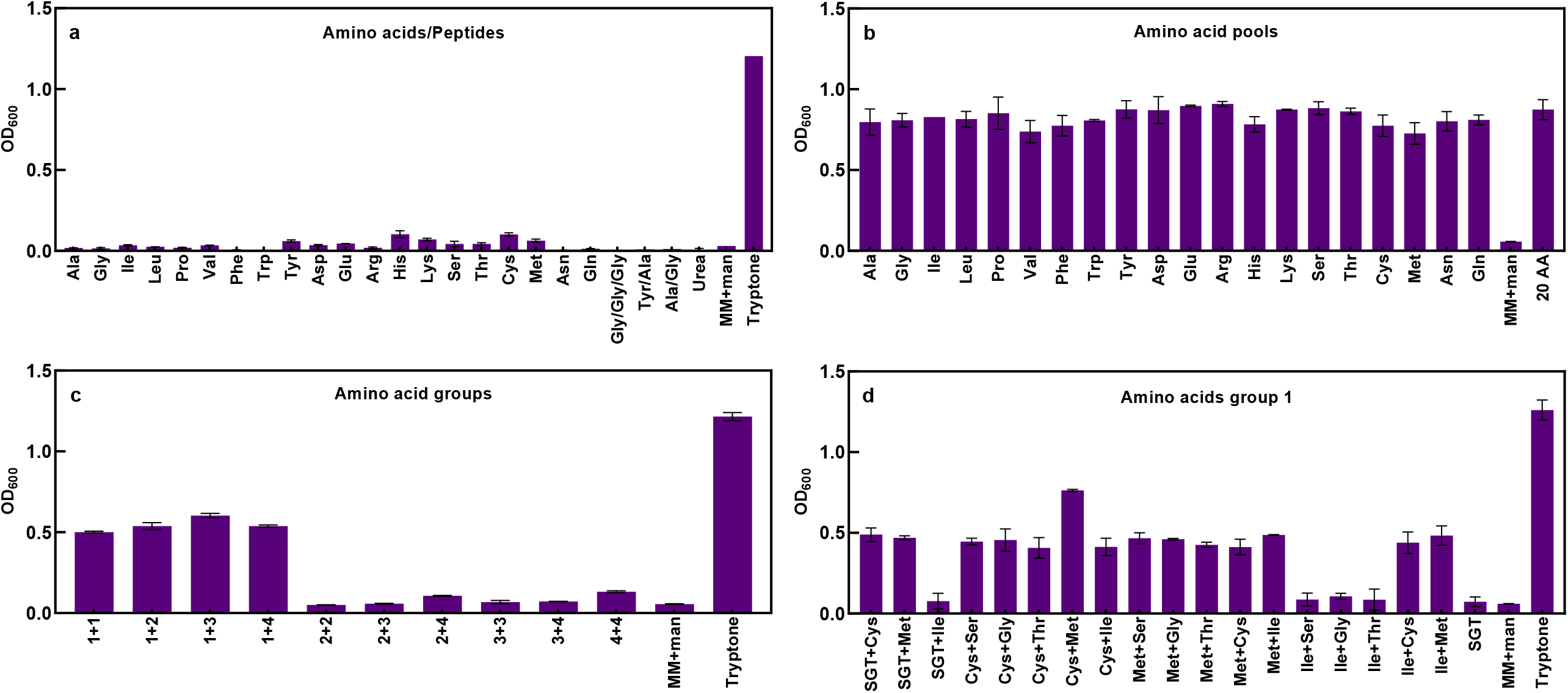
Amino acid growth requirement for *M. fakhimi.* MM supplemented with mannitol (5 g·L^-1^) (MM+man) was used as basal medium to dissolve the different amino acids/nutrients. The basal medium without any supplementation or supplemented with tryptone (1 g·L^-1^) were used as negative and positive controls, respectively. The pH was adjusted to 7.1±0.1 using KOH or HCl when needed and sterilized by 0.2 μm filters. The initial OD_600_ of *M. fakhimi* was 0.02, and the cultures were incubated at 25°C for 4 days before measuring final OD. **a) Growth on individual amino acids**. The basal medium was supplemented with Individual L-amino acids (8 mM, except for tyrosine, 2 mM), the indicated tri- and di-peptides (4 mM), or urea (2 g·L^-1^). **b) Growth on 19-containing amino acid pools**. The basal medium was supplemented with different pools of 19 L-amino acids (1 mM each, except for tyrosine, 0.1 mM). The pools are named with the amino acid that is missing in the pool. **c) Growth on amino acid groups**. Supplementation with different pools of L-amino acids (2 mM each amino acid, except for tyrosine, 0.2 mM). Pools were designed according to common biosynthetic pathways: Group 1: cysteine, methionine, glycine, threonine, isoleucine, serine; Group 2: aspartic acid, alanine, asparagine, lysine; Group 3: glutamic acid, glutamine, arginine, proline; Group 4: histidine, tryptophan, tyrosine, phenylalanine, leucine, valine. Combinations of different groups were assayed. **d) Growth on cysteine, methionine, glycine, threonine, isoleucine, and serine**. Supplementation with different combinations of the amino acids included in Group 1 (cysteine, methionine, glycine, threonine, isoleucine, and serine) (4 mM each amino acid, except for tyrosine, 1 mM). SGT: Serine, Glycine, Threonine.

Growth test using vitamins were assayed (**Fig. 4**). Importantly, for these growth studies, the bacterial precultures were done in MM supplemented with mannitol, cysteine, and methionine (and not in the LB medium). Partial restoration of the growth observed with tryptone was observed after the simultaneous supplementation with biotin and thiamine, and independently of cysteine and methionine supplementation (**Fig. 4**). It is possible that in the experiments depicted in Fig. 3, and opposite to what was shown on Fig. 4, the null effect of cysteine and methionine could be due to fact that the precultures were very rich in these two amino acids and inner reservoirs were still available for *M. fakhimi* sp. nov. cells. Further experiments with fast OD kinetics during 9 h (using similar conditions to Fig. 4) confirmed the auxotrophy for biotin and thiamine of *M. fakhimi* sp. nov. (**Supplemental Fig. 2**). Again, supplementation with cysteine and methionine had no effect. However, when the 20 essential amino acids were co-supplemented with the vitamins the maximal growth was observed. In this last scenario, it is possible that the amino acids were providing with extra C and N sources that improved the growth.

**Fig. 4.**
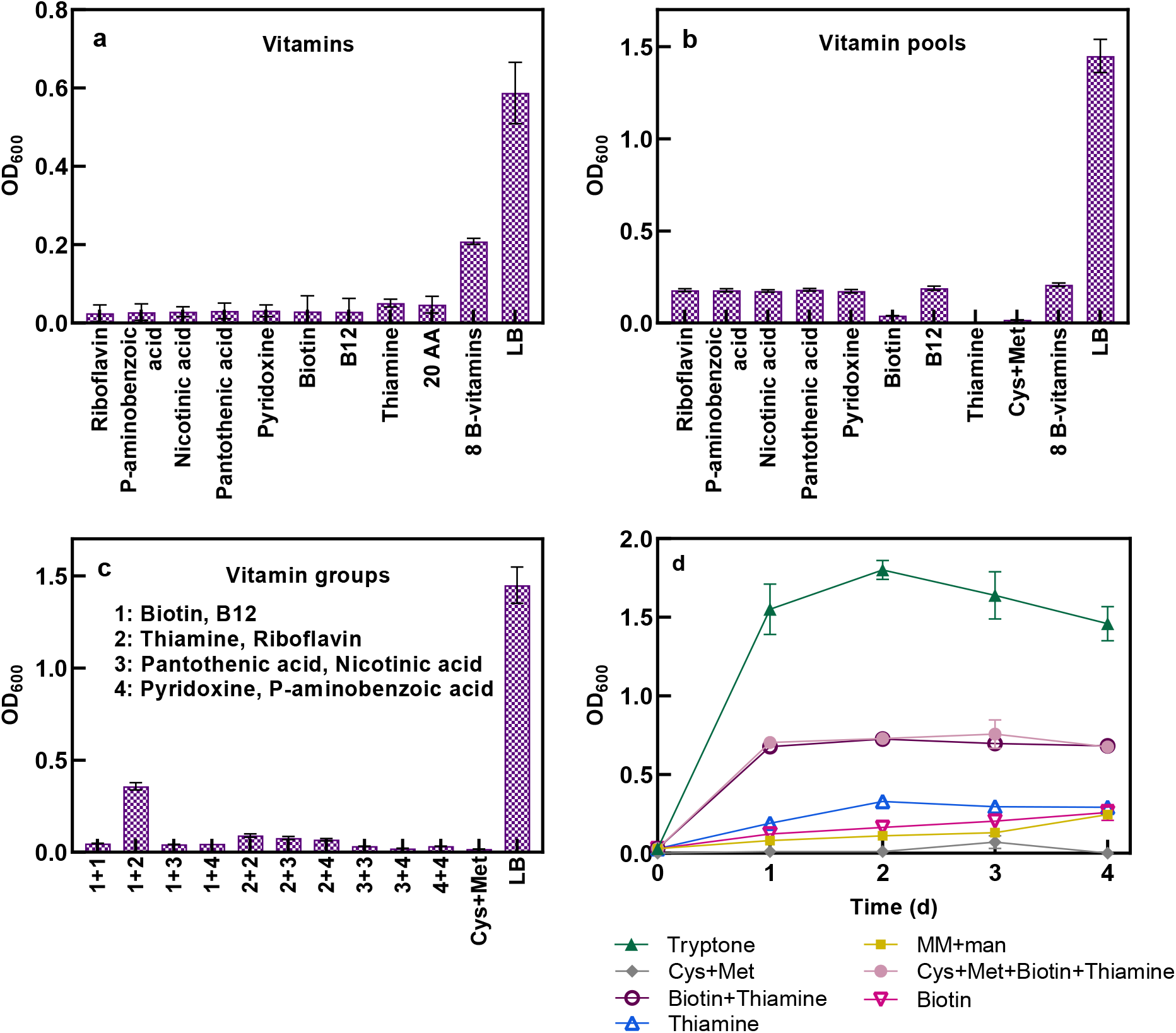
Vitamins growth requirements for *M. fakhimi sp. nov.* The initial inoculum of the bacterium was precultured twice in a row on MM supplemented with mannitol (5 g·L^-1^) cysteine (4mM) and methionine (4mM). MM supplemented with mannitol and 0.5 mM of all 20 amino was used as basal medium. MM with mannitol was used as negative control. All the cultures were incubated at 28° C. **a) Growth on vitamins**. The basal medium was supplemented with riboflavin (0.5 mg·L^-1^), p-aminobenzoic acid (0.1 m·g L^-1^), nicotinic acid (0.1 m·g L^-1^), pantothenic acid (0.1 m·g L^-1^), pyridoxine (0.1 m·g L^-1^), biotin (0.001 m·g L^-1^), vitamin B12 (0.001 m·g L^-1^), thiamine (0.001 m·g L^-1^) or all the 8 vitamins. These vitamins concentrations were used for all experiments. **b) Growth on vitamin pools**. <u>The basal medium</u> was supplemented with different pools of 7 vitamins. The pools are named with the vitamin that is missing in the pool. **c) Growth on vitamin groups**. <u>The basal medium</u> was supplemented with different pools of vitamins. Group 1: biotin and B12; Group 2: thiamine and riboflavin; Group 3: pantothenic acid and nicotinic acid; Group 4: pyridoxine and p-aminobenzoic acid. Binary combinations of these groups of vitamins were used. **d) Growth on different combinations of cysteine, methionine, biotin, and thiamine**. MM with mannitol was supplemented with the different nutrient combinations. MM with mannitol and tryptone was used as positive control, and MM with only mannitol was used as negative control.

To deconvolute the role of cysteine, methionine, biotin, and thiamine dependence during *M. fakhimi* sp. nov. growth, several consecutive culturing rounds in media with different combination of these nutrients were performed (**Fig. 5**). Only media containing the four nutrients allowed substantial growth after 5 consecutive culturing rounds, while the absence of either cysteine/methionine or thiamine/biotin resulted in very poor growth after two rounds (**Fig. 5**). Hence, *M. fakhimi* sp. nov. showed simultaneous growth dependence for methionine, cysteine, biotin, and thiamine.

**Fig. 5.**
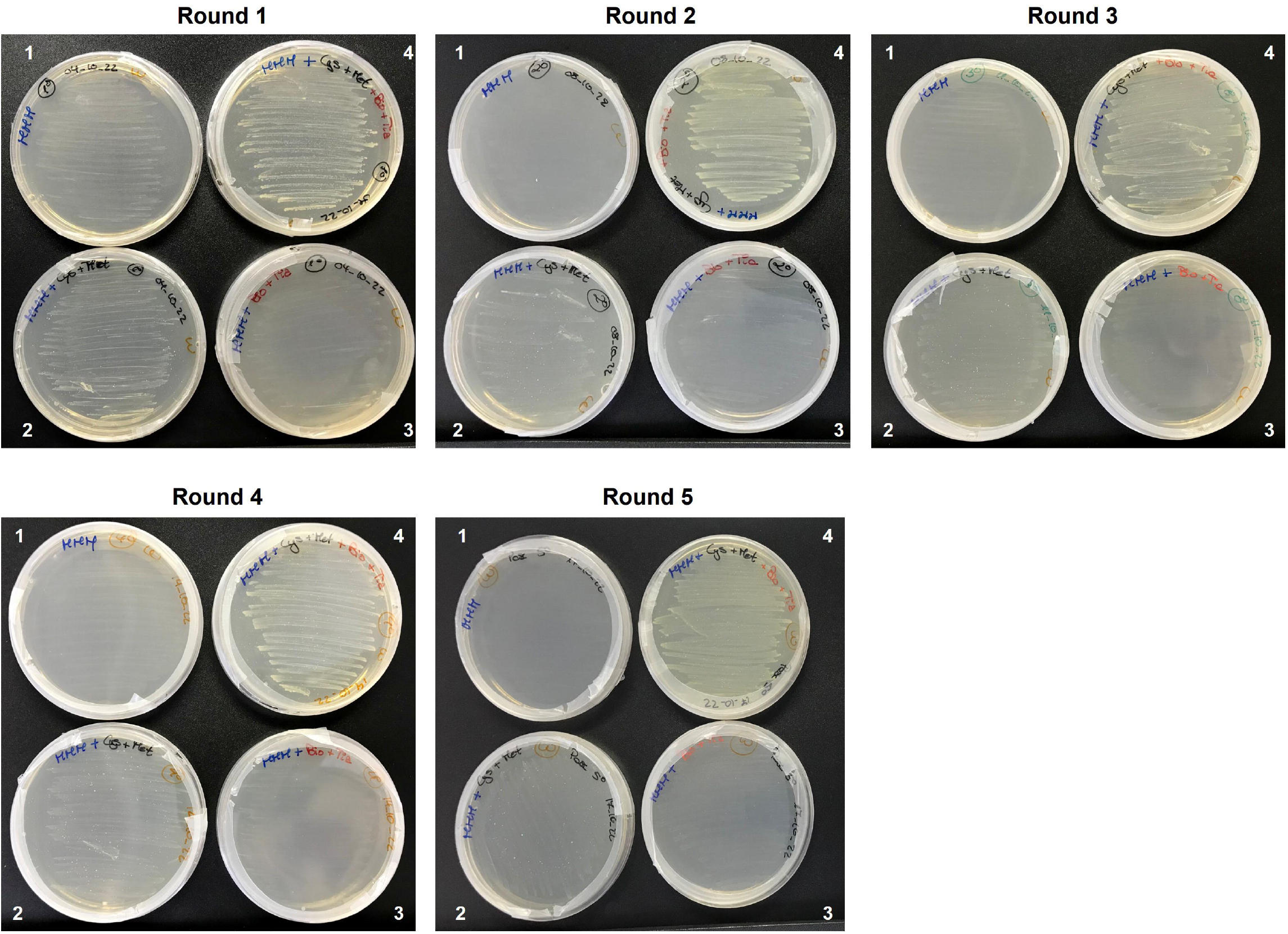
Consecutive plate streaking rounds in different media. *M. fakhimi* sp. nov. cells from an LB plate were streaked in plates without vitamin or amino acid supplementation (1), with cysteine and methionine (2), with biotin and thiamine (3), and with cysteine, methionine, biotin, and thiamine (4). All plates contained MM + mannitol (5 g·L^-1^). Plates were incubated at 28ºC for 3 days before, re-streaking on fresh plates with the corresponding media. Concentrations for amino acids and vitamins were of 4 mM and 0.001 mg·L^-1^, respectively.

According to the genome annotation and KEGG orthology assignments, *M. fakhimi* sp. nov. is deficient in the biosynthesis of both biotin and thiamine, which is in accordance with the experimental data. However, according to the KEGG orthology assignments, the biosynthetic pathways for cysteine and methionine biosynthesis are present in the *M. fakhimi* sp. nov. genome. Interestingly, there is not a single gene in the *M. fakhimi* sp. nov. genome for the sulfate assimilation pathway. There are not putative sulfate transporters, ATP sulfurylase, adenosine phosphosulfate kinase, adenosine phosphosulfate reductase or sulfite reductase genes. Consequently, *M. fakhimi* sp. nov. may not be able to reduce sulfate until 3’-phosphoadenylyl sulfate (PAPS) or sulfide, which are intermediates needed for the biosynthesis of many S-containing metabolites including cysteine and methionine (cysteine biosynthesis requires sulfide, and methionine biosynthesis requires cysteine). Note, that cysteine, methionine, biotin, and thiamine are sulfur S-containing metabolites. Hence, it is possible that the growth enhancement observed in Fig. 3 after cysteine and methionine supplementation is not directly related with cysteine or methionine auxotrophy, but indirectly by providing a reduced S source that can be used for the biosynthesis of S-metabolites.

The existence of two genes encoding for a putative sulfonate transport system (ssuA-, ssuB-, and ssuC-like) in the genome, suggests that this bacterium may uptake some S-reduced forms (Eichhorn et al., 2000; Kahnert et al., 2000; Endoh et al., 2003). Growth tests using some reduced forms of S (sulfite, sulfide, metabisulfite and thiosulfate) did not show growth restoration (**Supplemental Fig. 3a**). Additional growth tests using nucleosides/nucleotides also failed to restore growth (**Supplemental Fig. 3b**). Hence, these reduced forms of sulfur may not be used as S-source by bacteria or may be toxic at the concentrations used.

Extended growth test using media supplemented with methionine, cysteine, biotin and thiamine and different C sources were performed (**Table 2)**. *M. fakhimi* sp. nov. was able to efficiently grow on sucrose, maltose, many monosaccharides, and glycerol. Growth on lactose was poor. Little growth was observed with some organic acids (citric and lactic acids) and with five amino acids (alanine, arginine, glutamate, tyrosine, and glutamine). Efficient utilization of these five amino could probably provide both C and N sources. Growth was also evaluated in the LB medium at different temperatures and pHs **(Table 3)**. *M. fakhimi* sp. nov. had an optimal temperature growth range between 15-37<u>º</u>C, and in an optimal pH range between 6-9.5. No resistance to the antibiotics tetracycline, rifampicin, chloramphenicol and polymyxin (50 μg·mL^-1^ each) was observed (data not shown).

**Table 2.** Growth of *M. fakhimi* sp. nov. with different carbon sources. MM medium supplemented with cysteine (4 mM), methionine (4 mM), biotin (0.005 mg·L^-1^) and thiamin (0.005 mg·L^-1^) was used as basal medium. The different carbon sources and amino acids were added at 5 g·L^-1^ and 4 mM, respectively. ++, significant growth; +, poor growth; -, no growth

| Disaccharides |  | L-Amino acids |  |
| --- | --- | --- | --- |
| Sucrose | ++ | Alanine | + |
| Lactose | + | Glycine | - |
| Maltose | ++ | Isoleucine | - |
| Monosaccharides |  | Leucine | - |
| Glucose | ++ | Proline | - |
| Fructose | ++ | Valine | - |
| Galactose | ++ | Tryptophan | - |
| Mannitol | ++ | Lysine | - |
| Arabinose | ++ | Asparagine | - |
| Xylose | ++ | Methionine | - |
| Organic acids |  | Cysteine | - |
| Malic acid | - | Threonine | - |
| Tartaric acid | - | Histidine | - |
| n-butyric acid | - | Arginine | + |
| Formic acid | - | Glutamic acid | + |
| Acetic acid | - | Phenylalanine | - |
| Malonic acid | - | Tyrosine | + |
| Citric acid | + | Aspartic acid | - |
| Lactic acid | + | Serine | - |
| Other carbon sources |  | Glutamine | + |
| Glycerol | ++ |  |  |
| Ornithine | - |  |  |

**Table 3.** *M. fakhimi* sp. nov. growth at different temperatures and pH. LB medium was used in all the experiments. ++, significant growth; +, poor growth; -, no growth

| Temperature °C |  | pH |  |
| --- | --- | --- | --- |
| 10 | - | 4 | - |
| 15 | ++ | 5 | + |
| 20 | ++ | 6 | ++ |
| 24 | ++ | 7 | ++ |
| 30 | ++ | 8.5 | ++ |
| 37 | ++ | 9.5 | ++ |
| 42 | + | 10.5 | + |
| 45 | - |  |  |

Finally, *M. fakhimi* sp. nov. was cultivated under light or dark conditions to evaluate any potential induction of the yellow pigmentation. Light conditions slightly promoted pigmentation (**Supplemental Fig. 4a**). Pigments were extracted from a *M. fakhimi* sp. nov. culture and their wavelength spectrum were evaluated (**Supplemental Fig. 4b**). The typical previously described three peaks for decapornoxanthin at 413 nm, 437 nm, and 467 nm were observed (Mitra et al., 2021).

### Biotechnological importance of *Microbacterium fakhimi*

Microbacterium sp. is among the most common genera of bacterial endophytes and inhabitants of the rhizosphere (Rosenblueth and Martínez-Romero, 2006; Marquez-Santacruz et al., 2010b; Abadi et al., 2020). Microbacterium sp., is also frequently found within algal cultures (Burmølle et al., 2006; Jones et al., 2016), including, interestingly, *Chlamydomonas reinhardtii* cultures (Li et al., 2013; Mitra et al., 2021), which probably reflects frequent coexistence of *Chlamydomonas* sp. and *Microbacterium* sp. in natural ecosystems. Since M. *fakhimi* sp. nov. is unable to use sulfate as S source and has several auxotrophies such as for biotin and thiamine, its survival in natural ecosystems could be highly dependent on the surrounding microbial communities.

A recent publication has shown that the alga *Chlamydomonas reinhardtii* and *M. fakhimi* can stablish a mutualistic association when cocultured aerobically in the media containing mannitol and yeast extract (Fakhimi et al., 2023b). *M. fakhimi* sp. nov. provided the alga with acetic acid and ammonium (Fakhimi et al., 2023b), which promoted the alga growth, while the alga alleviated the bacterium auxotrophy for biotin and thiamine and its dependence for S-reduced sources. This association was mutually beneficial and prevented an excessive bacterial growth in cocultures, which could be one of the main drawbacks when algae-bacteria cocultures are used for biotechnological applications. In fact, the Chlamydomonas-*M. fakhimi* cocultures could led to excellent biomass production and bioremediation performance. Moreover, under hypoxic conditions *M. fakhimi* promoted a sustained H_2_ production of *C. reinhardtii*. H_2_ production was even possible in synthetic dairy wastewater (Fakhimi et al., 2023b). Interestingly, the algal H_2_ production and algal viability was even better when a multispecies association consisting in *C. reinhardtii, M. fakhimi* sp. nov., *Stenotrophomonas goyi* sp. nov. and *Bacillus cereus* was used (Fakhimi et al., 2023a; Fakhimi et al., 2023b). Using this multispecies consortium, the H_2_ production was concomitant with algal biomass generation. Despite *M. fakhimi* sp. nov. was the solely bacterium promoting H_2_ production, *S. goyi* sp. nov. and *B. cereus* played a role in benefiting algal viability under hypoxia. In this multispecies association, the metabolic interactions and its biotechnological interest needs to be better resolved.

## Author Contributions

Writing—original draft preparation, D.G.-B.; writing—review & editing, N.F.; Isolation of the *Microbacterium fakhimi* sp. nov., N.F.; design and execution of experiments, N.F., M.J.T, J.L.-D.; in silico analysis, D.G.-B.; results analysis and interpretation, N.F., D.G.-B. and A.D.; supervision, A.D. and D.G.-B.; project administration, A.G., E.F., A.D. and D.G.-B.; funding acquisition, A.G., E.F., A.D. and D.G.-B. All authors have read and agreed to the published version of the manuscript.

## Acknowledgments

The authors acknowledge Dr. Gregorio Galvéz Valdivieso for his unvaluable contribution to this research: perfection sometimes kills new discoveries. We thank the Bio knowledge Lab (BK-L) Ltd. for their kind support.

## Figure legends

**Supplemental Fig. 1. Plate with *Chlamydomonas reinhardtii* and *Microbacterium fakhimi* sp. nov. colonies**

**Supplemental Fig. 2. Fast growth kinetic for *M. fakhimi* sp. nov**. The initial inoculum of the bacterium was precultured twice in a row on MM supplemented with mannitol (5 g·L^-1^) (basal medium), cysteine (4mM) and methionine (4mM). The initial inoculum was set to OD 0.05, and the cultures were incubated at 30°C. **a)** MM with mannitol (basal medium) was considered as control. **b)** Basal medium was supplemented with cysteine and methionine, **c)** 20 essential amino acids, **d)** biotin and thiamine, **e)** cysteine, methionine, biotin, and thiamine, or **f)** 20 essential amino acids + biotin and thiamine. Cysteine and methionine were employed at 4 mM; biotin and thiamine at 0.005 mg·L^-1^; 20 amino acids at 1 mM each but tyrosine 0.5 mM.

**Supplemental Fig. 3. a) Nucleic acids, nucleosides and nucleotides growth requirement for *M. fakhimi sp. nov***. The initial bacterium inoculum was precultured two times in MM+5 g·L^-1^ mannitol + 0.5 mM of all 20 amino acids (basal medium) and then inoculated in the same (final OD 0.02) medium supplemented with 10 mg·L^-1^ (each) of purines (Xanthine, hypoxanthine, adenine and guanine), pyrimidines (uracil, thymine and cytosine), nucleosides (uridine, cytidine, xanthosine, inosine, adenosine, guanosine and thymidine) and nucleotides (uridine monophosphate, UMP; cytidine monophosphate, CMP; xanthosine monophosphate, XMP; adenosine monophosphate, AMP; guanosine monophosphate, GMP; and thymidine monophosphate, TMP). All the media were filter sterilized and cultures incubated at 28°C. Growth in the basal medium with and without mixture of all nucleic acids and their derivatives supplementation and LB was considere d as controls. **b) Reduced forms of S requirement for *M. fakhimi sp. nov***. Basal medium (MM+ 5 g·L^-1^ mannitol) was supplemented with different S reduced sources (0.125 g·L^-1^ each), including sodium sulfite, sodium sulfide hydrate, sodium metabisulfite and sodium thiosulfate. Basal medium without any supplementation and LB medium were used as negative and positive controls, respectively.

**Supplemental Fig. 4. a) Light dependence in *M. fakhimi* sp. nov. pigmentation. b) Absorption spectrum for extracted pigments**.

**Supplemental Table 1. Tentative genome annotation**

**Supplemental Table 2. Pairwise dDDH values between *M. fakhimi* sp. nov. and the closest type strains genomes**. The dDDH values are provided along with their confidence intervals (C.I.) for the three different GBDP formulas: a) formula d0 (a.k.a. GGDC formula 1): length of all HSPs divided by total genome length; b) formula d4 (a.k.a. GGDC formula 2): sum of all identities found in HSPs divided by overall HSP length; formula d6 (a.k.a. GGDC formula 3): sum of all identities found in HSPs divided by total genome length. Note: Formula d4 is independent of genome length and is thus robust against the use of incomplete draft genomes.

**Supplemental Table 3. Closest type strains**

**Supplemental Table 4. BlastKOALA KEGG orthology assignments of** *M. fakhimi* **putative proteins**

