## Supplementary figures and images for "*Draft genome sequence of Microbacterium fakhimi* sp. nov., a novel bacterium associated with the alga *Chlamydomonas reinhardtii*"

### Supplemental Figure 1

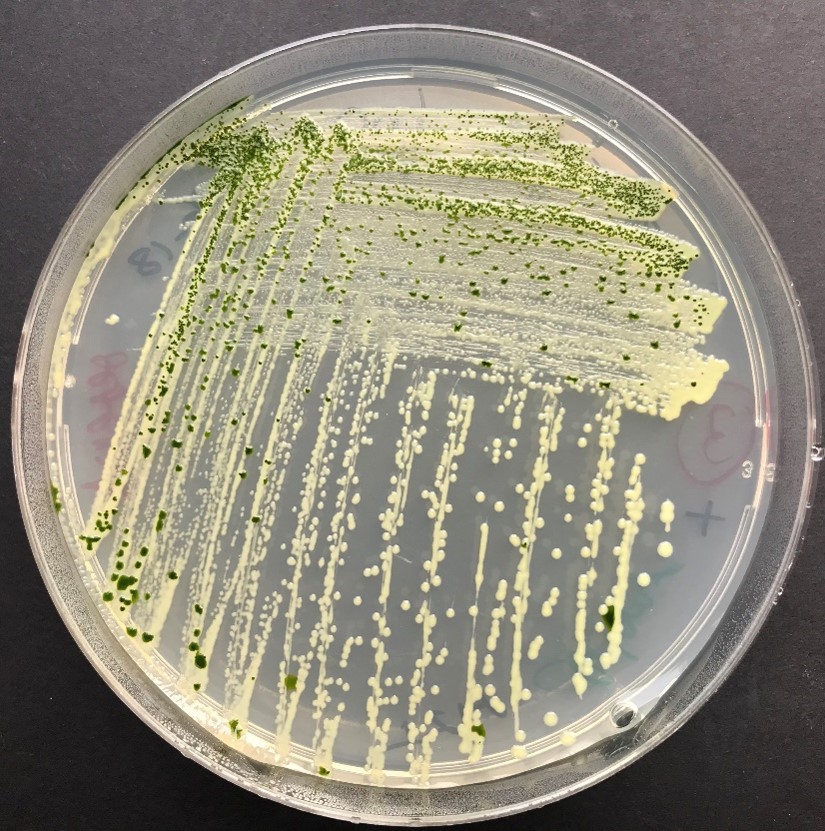

### Supplemental Figure 2

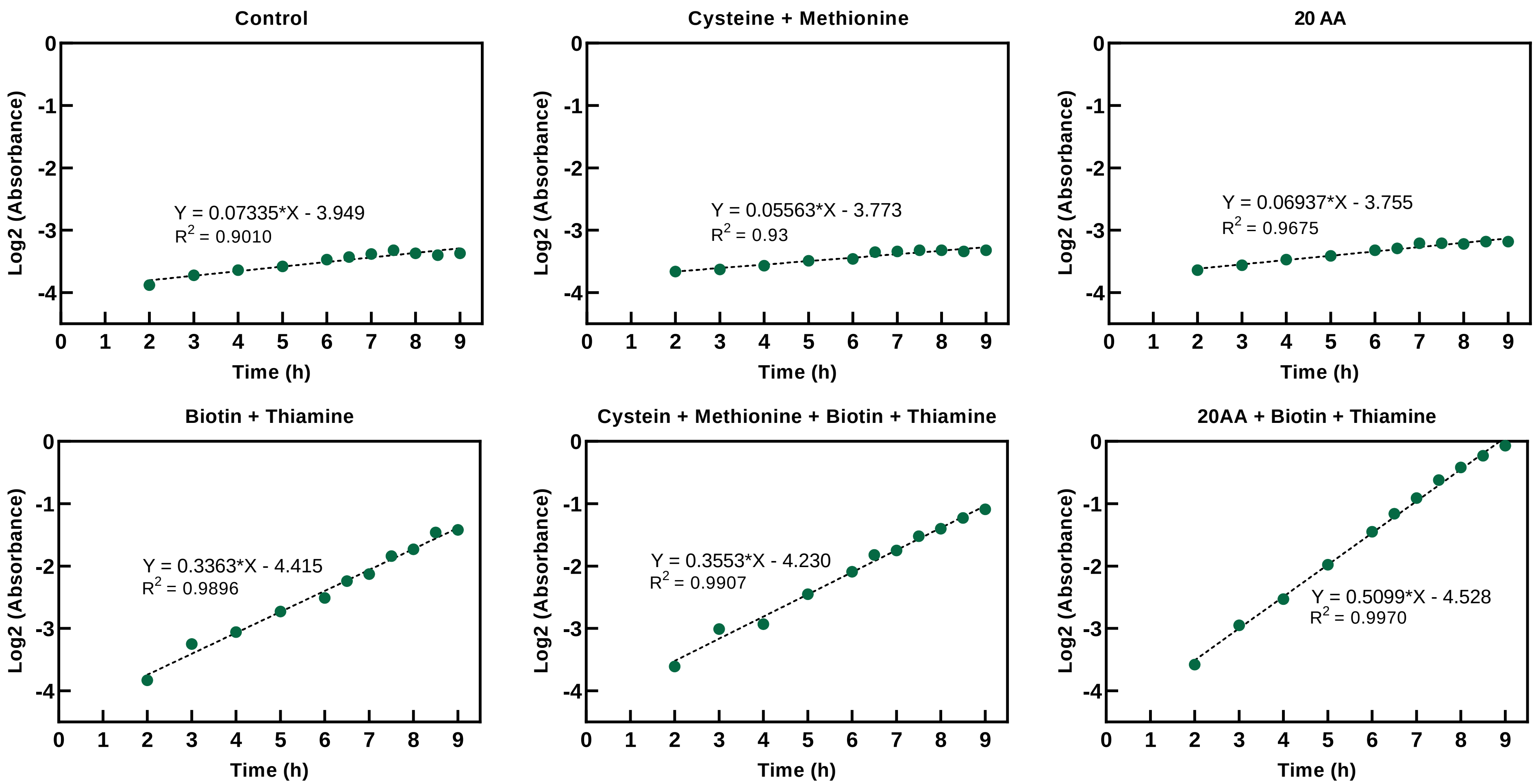

### Supplemental Figure 3

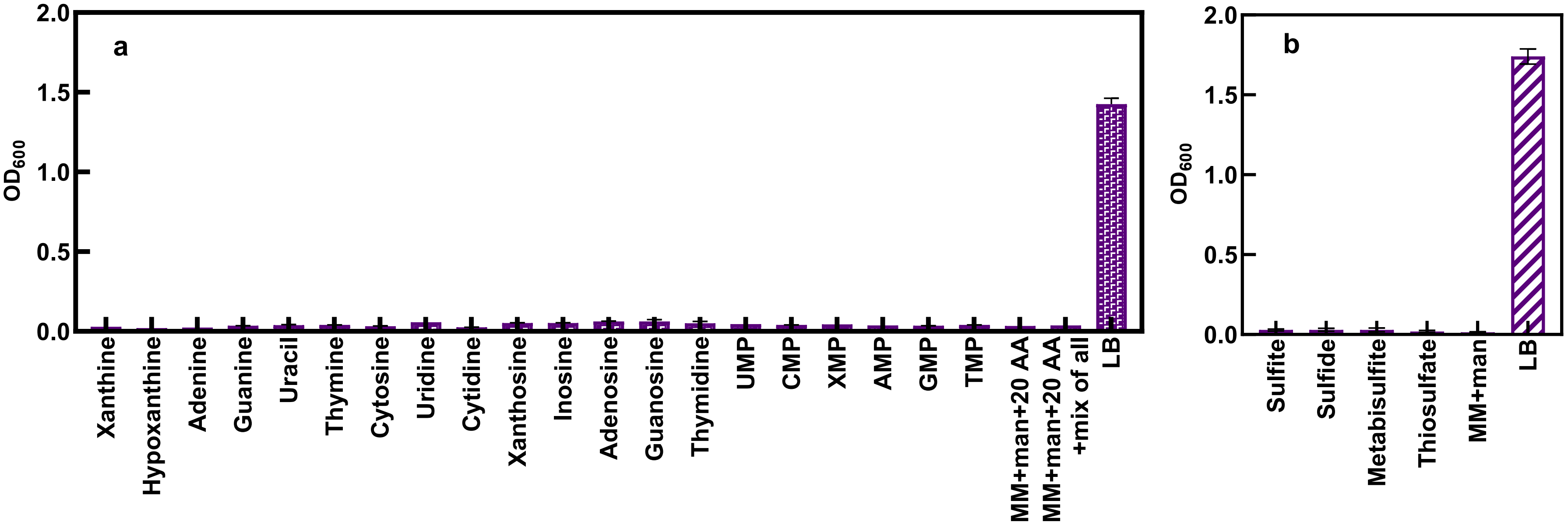

### Supplemental Figure 4

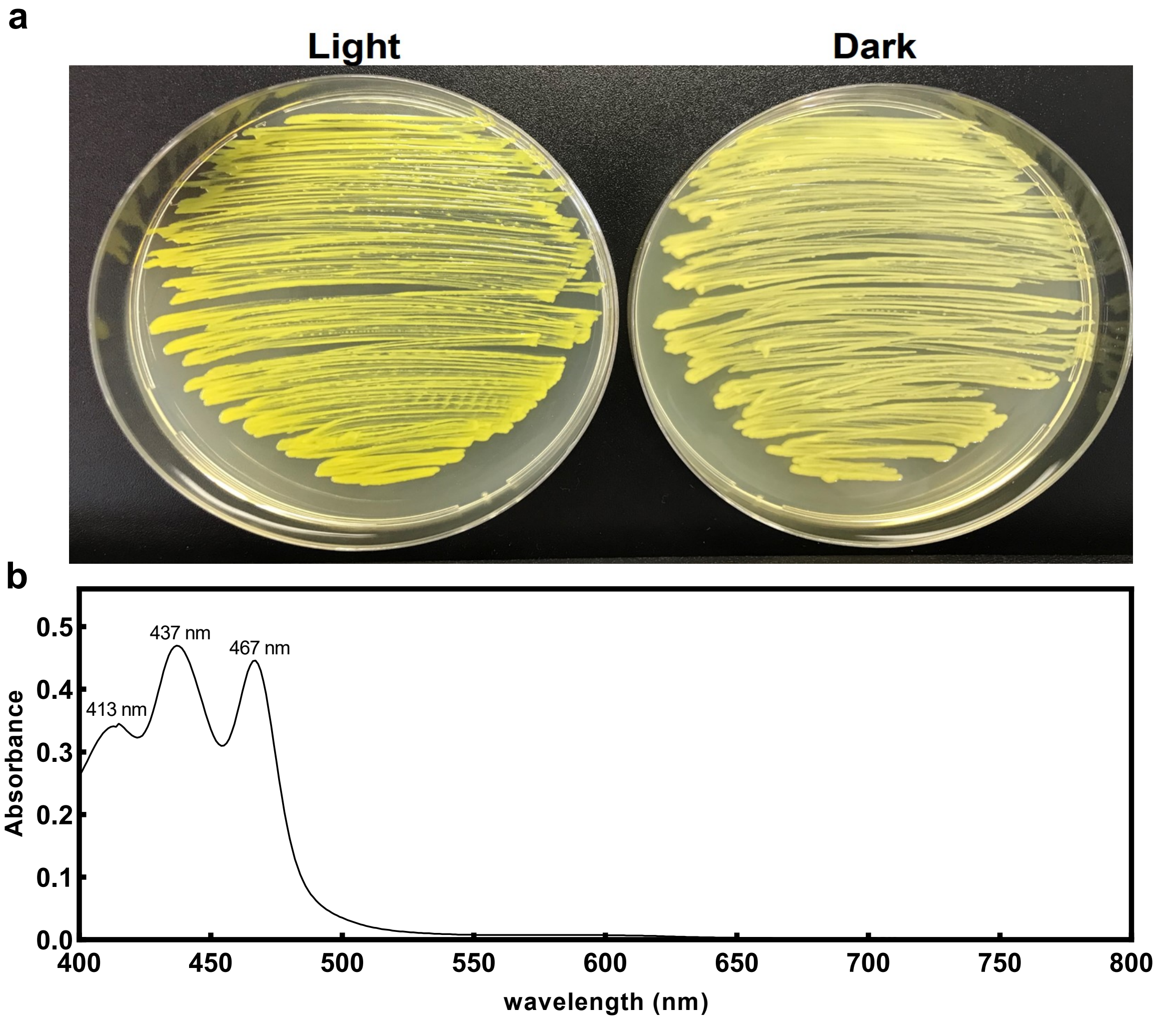
