## Supplemental Table 2 for "*Draft genome sequence of Microbacterium fakhimi* sp. nov., a novel bacterium associated with the alga *Chlamydomonas reinhardtii*"

| **Closest strains to *Microbacterium fakhimi*** | **dDDH**  **(d0, in %)** | **C.I.**  **(d0, in %)** | **dDDH**  **(d4, in %)** | **C.I.**  **(d4, in %)** | **dDDH**  **(d6, in %)** | **C.I.**  **(d6, in %)** | **G+C content difference (in %)** |
| --- | --- | --- | --- | --- | --- | --- | --- |
| *Microbacterium phyllosphaerae DSM 13468* | 61,9 | [58.2 - 65.5] | 30,8 | [28.4 - 33.3] | 53,1 | [49.9 - 56.1] | 0,19 |
| *Microbacterium hydrocarbonoxydans DSM 16089* | 58,4 | [54.8 - 62.0] | 28,9 | [26.6 - 31.4] | 49,6 | [46.5 - 52.6] | 0,67 |
| *Microbacterium foliorum DSM 12966* | 50,7 | [47.2 - 54.1] | 27,7 | [25.3 - 30.2] | 43,6 | [40.6 - 46.6] | 0,79 |
| *Microbacterium algeriense G1T* | 39,5 | [36.2 - 43.0] | 24,6 | [22.3 - 27.1] | 34,6 | [31.6 - 37.7] | 1,69 |
| *Microbacterium saperdae DSM 20169* | 35,1 | [31.7 - 38.6] | 24 | [21.7 - 26.4] | 31,2 | [28.3 - 34.3] | 0,38 |
| *Microbacterium maritypicum DSM 12512* | 37,7 | [34.3 - 41.2] | 23,9 | [21.6 - 26.4] | 33 | [30.1 - 36.1] | 0,56 |
| *Microbacterium saperdae JCM 1352* | 34,6 | [31.3 - 38.2] | 23,9 | [21.6 - 26.4] | 30,9 | [28.0 - 34.0] | 0,2 |
| *Microbacterium liquefaciens JCM 3879* | 35,1 | [31.7 - 38.6] | 23,8 | [21.5 - 26.3] | 31,2 | [28.2 - 34.3] | 0,27 |
| *Microbacterium liquefaciens NBRC 15037* | 35,1 | [31.7 - 38.6] | 23,8 | [21.5 - 26.3] | 31,2 | [28.3 - 34.3] | 0,32 |
| *Microbacterium paraoxydans DSM 15019* | 34,1 | [30.7 - 37.6] | 23,6 | [21.3 - 26.1] | 30,4 | [27.5 - 33.5] | 2,11 |
| *Microbacterium oxydans NBRC 15586* | 37,3 | [33.9 - 40.8] | 23,2 | [20.9 - 25.7] | 32,5 | [29.6 - 35.6] | 0,37 |
| *Microbacterium oxydans DSM 20578* | 37,3 | [34.0 - 40.8] | 23,2 | [20.9 - 25.7] | 32,5 | [29.6 - 35.6] | 0,35 |

**Supplemental Table 2. Pairwise dDDH values between M. fakhimi sp. nov and the closest type strains genomes**. The dDDH values are provided along with their confidence intervals (C.I.) for the three different GBDP formulas: a) formula d0 (a.k.a. GGDC formula 1): length of all HSPs divided by total genome length; b) formula d4 (a.k.a. GGDC formula 2): sum of all identities found in HSPs divided by overall HSP length; formula d6 (a.k.a. GGDC formula 3): sum of all identities found in HSPs divided by total genome length. Note: Formula d4 is independent of genome length and is thus robust against the use of incomplete draft genomes.
